# SINCERITIES: Inferring gene regulatory networks from time-stamped single cell transcriptional expression profiles

**DOI:** 10.1101/089110

**Authors:** Nan Papili Gao, S.M. Minhaz Ud-Dean, Rudiyanto Gunawan

**Affiliations:** Institute for Chemical and Bioengineering, ETH Zurich, Zurich, Switzerland; Swiss Institute of Bioinformatics, Lausanne, Switzerland; Werner Siemens Imaging Center, Department of Preclinical Imaging and Radiopharmacy, Eberhard Karls University Tuebingen, Tuebingen, Germany

**Keywords:** network inference, single cell, gene expression, gene regulatory network, time-stamped cross-sectional data

## Abstract

Recent advances in single cell transcriptional profiling open up a new avenue in studying the functional role of cell-to-cell variability in physiological processes such as stem cell differentiation. In this work, we developed a novel algorithm called SINCERITIES (SINgle CEll Regularized Inference using TIme-stamped Expression profileS), for the inference of gene regulatory networks (GRNs) from single cell transcriptional expression data. In particular, we focused on time-stamped cross-sectional expression data, a common type of dataset generated from transcriptional profiling of single cells collected at multiple time points after cell stimulation. SINCERITIES recovers the regulatory (causal) relationships among genes by employing regularized linear regression, particularly ridge regression, using temporal changes in the distributions of gene expressions. Meanwhile, the modes of the gene regulations (activation and repression) come from partial correlation analyses between pairs of genes. We demonstrated the efficacy of SINCERITIES in inferring GRNs using simulated time-stamped *in silico* single cell expression data and single transcriptional profiling of THP-1 monocytic human leukemia cell differentiation. The case studies showed that SINCERITIES could provide accurate GRN predictions, significantly better than other GRN inference algorithms such as TSNI, GENIE3 and JUMP3. Meanwhile, SINCERITIES has a low computational complexity and is amenable to problems of extremely large dimensionality.

## Background

Cell profiling technologies have enabled scientists to measure intracellular molecules (DNA, RNA, proteins, metabolites) at whole-genome level and down to single cell resolution. Over the last decade, high-throughput single cell assays have experienced tremendous progress, thanks to advanced microfluidics techniques and increased sensitivity in cell profiling assays. For example, the Fluidigm Dynamic Array platform employs integrated fluidics circuitry to capture single cells (up to 96 cells per run) for transcriptional expression profiling using quantitative RT-PCR (qRT-PCR) or RNA-sequencing (RNA-seq) [1]. The ability to assay individual cells and to examine intra-population cellular heterogeneity brings great benefits to fields such as stem cell and cancer biology. In the last few years, single cell analyses have demonstrated the ubiquity of cellular heterogeneity, even within cell populations or cell types that have been traditionally perceived as homogeneous [2–6]. Meanwhile, many single cell studies have provided evidence for the physiological roles of cell-to-cell variability in normal and diseased cells [7–11].

Single cell transcriptional profiling overcomes many issues associated with population-average or bulk data that mask cellular heterogeneity (e.g. Simpson’s paradox [12]), thereby presenting new means for understanding biology. The number of bioinformatics tools for analyzing single cell expression data has proliferated in recent years [13–15]. A class of these algorithms concerns with the deconvolution of cell populations and tissues to elucidate population substructures and identify known and novel cell subtypes [16–20]. These algorithms often apply or modify existing clustering and dimensionality reduction algorithms, such as PCA, tSNE and diffusion maps, to accommodate single cell data. Another class of algorithms deals with the ordering of cells within the cell population along a perceived transition path between different cell states (e.g. Monocle [21], Wanderlust [22], SCUBA [23] and TSCAN [24]). Such cell ordering produces a trajectory in the state space of gene expression corresponding to a physiological transition, such as stem cell differentiation process.

The third class of algorithms considers gene regulatory network (GRN) inference. A GRN is a network graph, where the nodes of this graph represent genes and the edges represent gene-gene interactions. The most common gene networks created from single cell transcriptional data have undirected edges (see for example [11,25,26]), where such edges indicate associations among genes, for example co-expression or co-regulation relationships. In contrast, the focus of our work is inferring GRNs with directed (causal) edges, where an edge pointing from gene *i* to gene *j* implies that the protein product(s) of gene *i* regulates the expression of gene *j* (e.g., gene *i* encodes a transcription factor of gene *j*). The edges may also have signs, representing the modes of the gene regulation: positive for activation and negative for repression. In comparison to the other two classes of algorithms, there have been lesser algorithmic developments on the inference of such GRNs from single cell transcriptional profiles, possibly because of the extreme difficulty in this task [13–15].

One of the challenges in using single cell expression data for GRN inference is the high data dropout rate due to transcriptional bursting of gene expression process, leading to zero-inflated dataset [13–15]. In addition, single cell profiling techniques such as qRT-PCR and RNA-seq use cell lysates, and consequently, the identities of the cells are lost. The resulting data therefore provide only cross-sectional information of the cell population. A few GRN inference methods have previously been proposed based on Boolean network model [27–29], stochastic gene expression model [30], and a combination of machine learning and nonlinear differential equation model [31]. However, none of these methods use time point information directly in the GRN inference. In general, temporal data possess more information than static or single time-point data, especially for the determination of causal networks [32]. For these reasons, here we consider time-stamped cross-sectional single cell transcriptional profiles, i.e. the expression profiles of single cells taken at multiple time points after cell stimulation. Such type of dataset is commonly generated in studies of cell differentiation process, where stem cells are induced to differentiate at the beginning of the experiment and afterwards cells are collected at multiple time points for single cell analysis [11,25,33].

In this work, we created a network inference algorithm, called SINCERITIES (SINgle CEll Regularized Inference using TIme-stamped Expression profileS). The GRN inference was formulated as regularized linear regressions based on temporal changes of the gene expression distributions. The modes of the gene regulations, i.e the signs of the edges, were determined using partial correlation analyses. We demonstrated the efficacy of SINCERITIES using *in silico* time-stamped single cell expression profiles, as well as time-stamped cross-sectional transcriptional profiles of THP-1 human myeloid monocytic leukemia cells [25]. We also compared SINCERITIES to existing GRN inference algorithms developed for time series expression data, namely TSNI [34] and JUMP3 [35], and to a tree-based GRN inference algorithm GENIE3 [36]. The case studies illustrated the efficacy of SINCERITIES in extracting accurate GRN, by taking advantage of temporal information in time-stamped single cell expression data.

## Results

### Gene regulatory network inference using SINCERITIES

Below, we provide a brief description of SINCERITIES. More details of SINCERITIES can be found in Methods. In the following, let *m* be the number of genes, *n* be the number of measurement time points, and *sk* be the number of cells in the *k*-th time point sample (*k* = 1, 2, …, *n*). Figure 1 illustrates the main steps of SINCERITIES. The time-stamped cross-sectional dataset (see Fig. 1A) comprises *n* data matrices *E*_*sk* *xm*_,where the matrix element *E*_*ik,j*_ is the transcriptional expression value of gene *j*, i.e. the amount of mRNA molecules of gene *j* in the *i*-th cell at the *k-*th time point. SINCERITIES is based on the assumption that changes in the expression of a transcription factor will alter the expression of the target genes. Thus, in the first step of SINCERITIES (see Fig. 1B), we quantify the temporal changes in the expression of each individual gene by computing the distance of the marginal gene expression distributions between two subsequent time points. For the distributional distance (DD) metric, we use the Kolmogorov-Smirnov (KS) distance, i.e. the maximum absolute difference between two cumulative density functions (see Methods) [37]. By using a DD metric, we could account for not only the overall changes in the gene expression distribution, but also the shift in the fractions of dropouts (i.e. genes with zero mRNA count). The performance of SINCERITIES did not depend sensitively on the choice of the DD metric (see Methods). Since the time windows may not necessarily be uniform, the DD values are normalized by the time step size.

**Figure 1.**
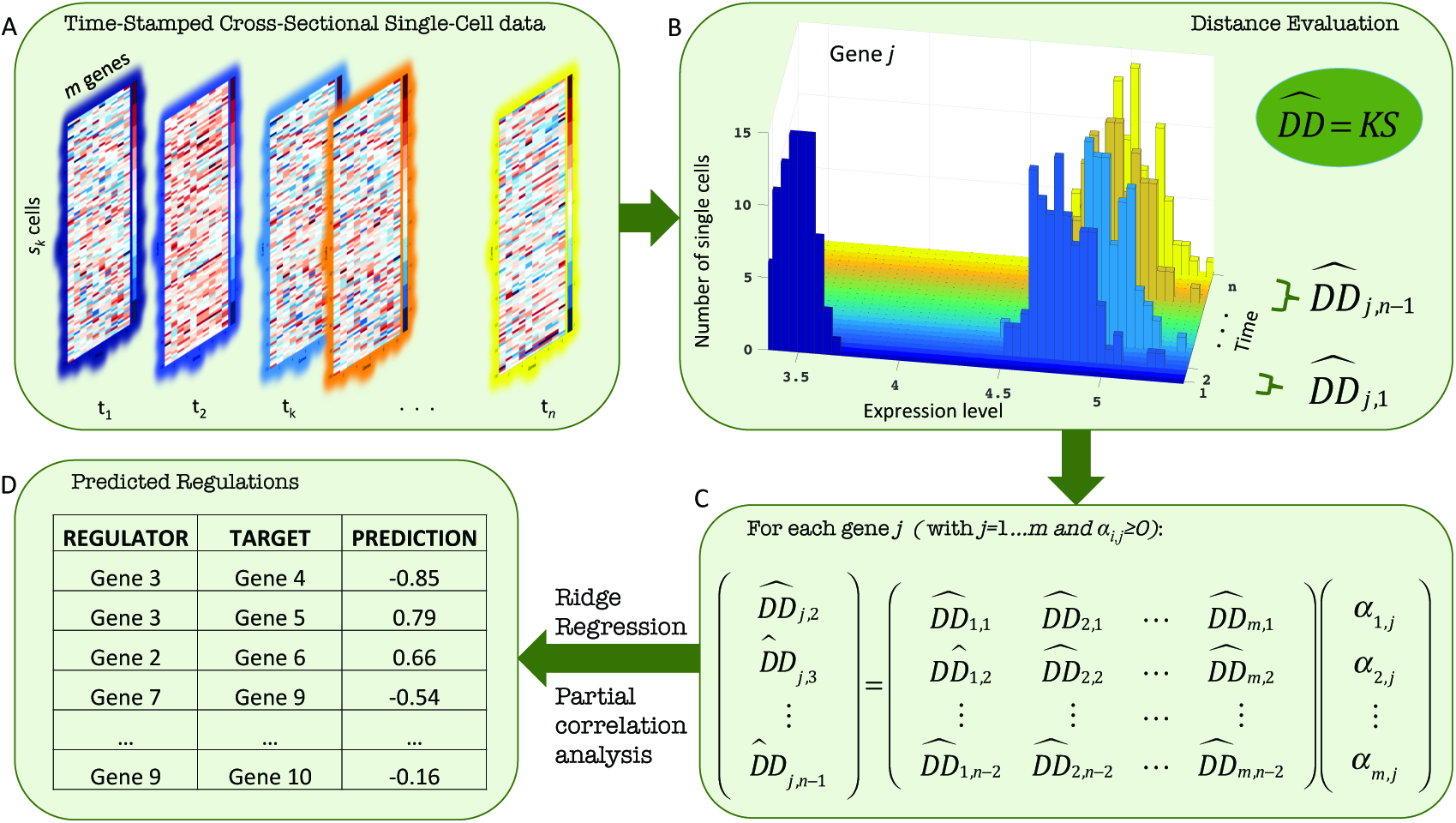
The workflow of SINCERITIES. (A) Input: time-stamped cross-sectional data of gene expression. (B) Step 1: calculation of normalised distribution distance of gene expression distributions over each time step; (C) Step 2: formulation of the GRN inference as a linear regression problem; (D) Output: edge predictions of the GRN.

In order to establish directed (causal) edges in the GRN, we treat the change in the expression of a transcription factor in a given time window as a perturbation. Furthermore, we assume that such a perturbation will cause a proportional shift in the gene expression distributions of the corresponding target genes in the next time window. As shown in Fig. 1C, the GRN inference in SINCERITIES involves solving *m* independent linear regressions. More specifically, for each gene *j*, we formulate a linear regression using the normalized DDs of this gene at time windows *l*+1, denoted by 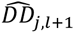 (I = 1,2,…, *n* — 1), as the response (dependent) variable, while setting the normalized DDs of all other genes at the previous time window *l* 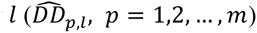 as the regressor (independent) variables.

The linear regression is thus given by:

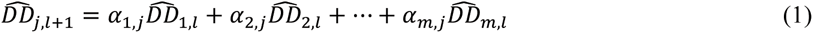

where *α*_*p,j*_ is the regression coefficient describing the influence of gene *p* on gene *j*. The least square solution vector *α*^*\**^_*j*_ is constrained to be non-negative since DDs take only non-negative values. The linear regression above is often underdetermined as the number of genes typically exceeds the number of time windows. For this reason, we employ a penalized least square approach to obtain *α*^*^_*j*_ using an L_2_-norm penalty, also known as ridge regression or Tikhonov regularization (see Methods for more details). SINCERITIES relies on GLMNET [38] to compute the solution vector *α*^*^ for each gene *j*, using leave-one-out cross-validation (LOOCV) for determining the weight of the penalty term. Upon completion, SINCERITIES produces a ranked list of all possible edges in the GRN (a total of *m*^2^ edges)in descending order of *α*_*p,j*_ values (see Fig. 1D). Larger *α*_*p,j*_ indicates higher confidence that the corresponding edge exists (i.e. the edge *p → j*). For the mode (sign) of the gene regulatory edges, SINCERITIES uses partial correlation analyses on the expressions of every gene pair, controlling for the other genes (see Methods). The sign of an edge is set to the sign of the corresponding partial correlation. In other words, a positive (negative) correlation is taken as an indication of activation (repression).

Presently, SINCERITIES cannot directly handle single cell data from stem cell differentiation process that produces more than one cell type (i.e. branching). In such a scenario, a pre-processing step is needed to group cells into individual cell lineages (for example, using time-variant clustering [39]), and SINCERITIES could subsequently be applied to data from each differentiation branch. In the case studies, we tested SINCERITIES performance in inferring moderately sized GRNs (<50 genes). While there exist no technical limitation in applying SINCERITIES to single cell expression data with many ore genes, for example RNA-seq data, we expect that network inferability issue would become important in such an inference [40,41]. Finally, SINCERITIES currently could not accommodate datasets with fewer than five time points due to the limitation of LOOCV.

### Evaluation of SINCERITIES on *in silico* single cell data

To evaluate the efficacy of SINCERITIES, we simulated *in silico* time-stamped single cell expression datasets using 10-gene and 20-gene gold standard GRNs. The gold standard GRNs comprise 40 random subnetworks of *Escherichia coli* and *Saccharomyces cerevisiae* GRNs, i.e. ten networks for each size and from each species (see Additional file 1), generated using GeneNetWeaver [42]. For the main dataset, we simulated single cell gene expression data for 100 cells at 8 unevenly spaced time points using a stochastic differential equation model (see Methods). In order to test the robustness of SINCERITIES with respect to the number of sampling time points and to the degree of stochasticity in the gene expression, we further generated supplementary datasets using the 10-gene GRNs above, for varying degrees of intrinsic noise (by changing *σ* parameter, see Methods) and different numbers of sampling time points. In the gold standard GRNs, we assumed that there exist no self-regulatory edges, since some of the existing algorithms used in the comparison, namely GENIE3 and JUMP3, do not identify or remove such edges from GRN predictions.

We assessed the performance of SINCERITIES by evaluating the areas under the Receiver Operating Characteristic (AUROC) and the Precision-Recall curve (AUPR) [43]. Higher AUROC and AUPR values indicate more accurate GRN predictions. For this purpose, we computed the numbers of true positive (TP), true negative (TN), false positive (FP) and false negative (FN) edges by comparing the regulatory edges in the gold standard network with the top *q* edges from the ranked list output of SINCERITIES. When considering GRNs with signed edges, a true positive prediction referred to the correct prediction of an edge and its sign. The ROC curve was constructed by plotting the true positive rates (TPR = TP/(TP+FN)) versus the false positive rates (FPR = FP/(FP+TN)) for increasing *q* (*q* = 1,2, …, *m*^2^). Similarly, the precision (TP/(TP+FP)) and recall (TP/(TP+FN)) curve was plotted for increasing *q*.

Table 1 gives the AUROC and AUPR values of SINCERITIES predictions for the main dataset, respecting the sign of the gene regulatory edges. As expected, the larger (20-gene) GRNs had lower AUROC and AUPR values, indicating that they were more difficult to infer than the smaller (10-gene) GRNs. Meanwhile, Table 2 shows the mean AUROC and AUPR values of SINCERITIES for the supplementary dataset. In general, the performance of SINCERITIES decreased slightly with increasing intrinsic stochasticity. Meanwhile, decreasing the number of time points did not appreciably change the performance of SINCERITIES.

**Table 1.** Performance of SINCERITIES in the inference of gold standard GRNs.

|  | SINCERITIES |  |  |  |
| --- | --- | --- | --- | --- |
|  | 10-GENE NETWORK |  | 20-GENE NETWORK |  |
|  | AUROC | AUPR | AUROC | AUPR |
| Network E. coli 1 | 0.65 | 0.14 | 0.48 | 0.09 |
| Network E. coli 2 | 0.71 | 0.15 | 0.44 | 0.06 |
| Network E. coli 3 | 0.77 | 0.15 | 0.75 | 0.19 |
| Network E. coli 4 | 0.85 | 0.36 | 0.57 | 0.08 |
| Network E. coli 5 | 0.80 | 0.19 | 0.55 | 0.07 |
| Network E. coli 6 | 0.59 | 0.12 | 0.81 | 0.27 |
| Network E. coli 7 | 0.54 | 0.17 | 0.75 | 0.16 |
| Network E. coli 8 | 0.83 | 0.23 | 0.82 | 0.28 |
| Network E. coli 9 | 0.79 | 0.29 | 0.71 | 0.10 |
| Network E. coli 10 | 0.88 | 0.35 | 0.58 | 0.07 |
| Network Yeast 11 | 0.69 | 0.26 | 0.70 | 0.18 |
| Network Yeast 12 | 0.64 | 0.13 | 0.73 | 0.27 |
| Network Yeast 13 | 0.84 | 0.68 | 0.60 | 0.06 |
| Network Yeast 14 | 0.84 | 0.57 | 0.63 | 0.09 |
| Network Yeast 15 | 0.86 | 0.44 | 0.71 | 0.31 |
| Network Yeast 16 | 0.90 | 0.48 | 0.70 | 0.17 |
| Network Yeast 17 | 0.78 | 0.39 | 0.73 | 0.13 |
| Network Yeast 18 | 0.92 | 0.72 | 0.65 | 0.17 |
| Network Yeast 19 | 0.73 | 0.23 | 0.78 | 0.26 |
| Network Yeast 20 | 0.94 | 0.73 | 0.73 | 0.20 |
| **Mean** | **0.78** | **0.34** | **0.67** | **0.16** |
| **± SD** | **0.11** | **0.20** | **0.10** | **0.08** |

**Table 2.**
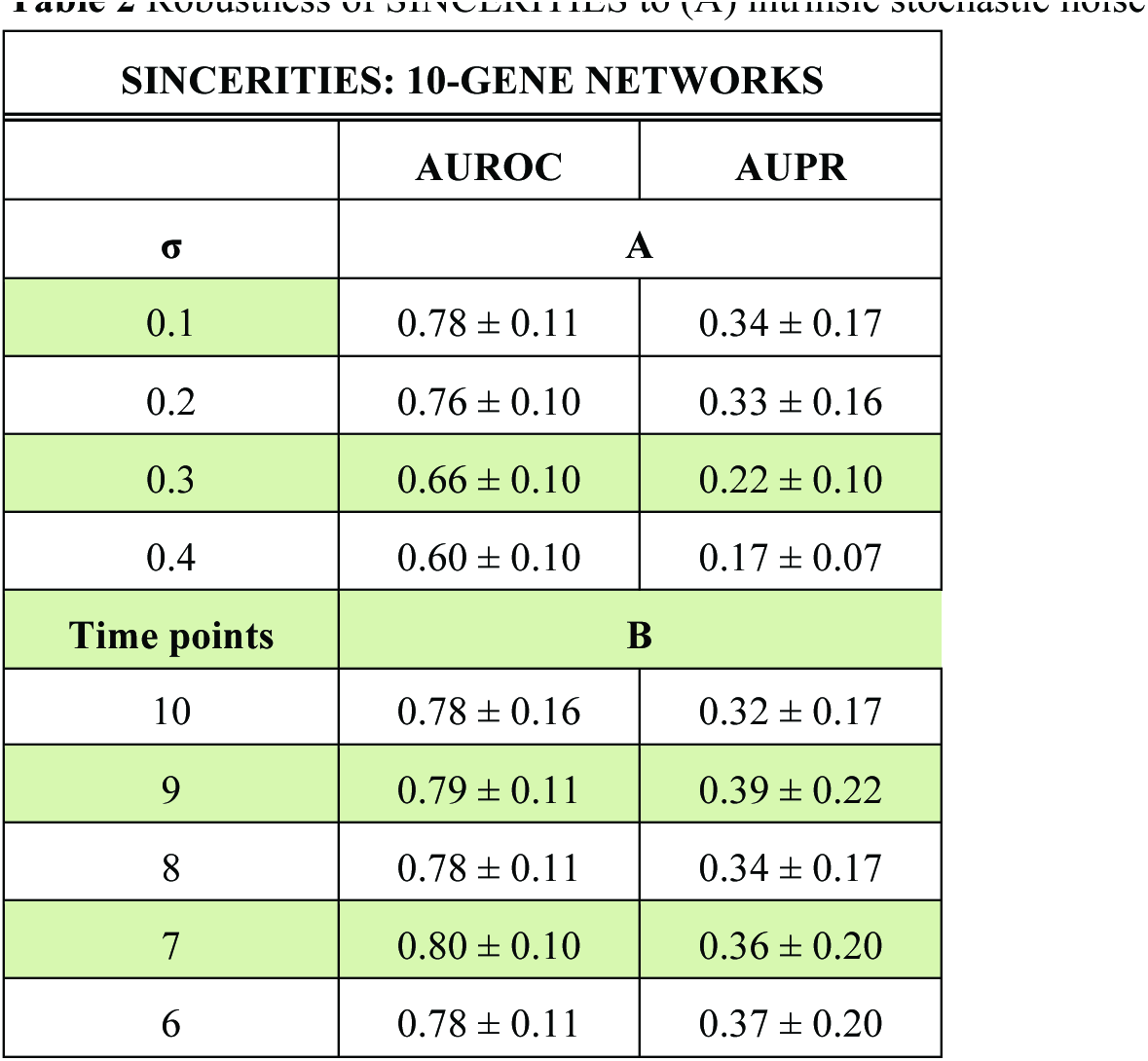
Robustness of SINCERITIES to (A) intrinsic stochastic noise and (B) number of time points.

| **SINCERITIES: 10-GENE NETWORKS** |  |  |
| --- | --- | --- |
|  | **AUROC** | **AUPR** |
| **$\sigma$** | **A** |  |
| 0.1 | $0.78 \pm 0.11$ | $0.34 \pm 0.17$ |
| 0.2 | $0.76 \pm 0.10$ | $0.33 \pm 0.16$ |
| 0.3 | $0.66 \pm 0.10$ | $0.22 \pm 0.10$ |
| 0.4 | $0.60 \pm 0.10$ | $0.17 \pm 0.07$ |
| **Time points** | **B** |  |
| 10 | $0.78 \pm 0.16$ | $0.32 \pm 0.17$ |
| 9 | $0.79 \pm 0.11$ | $0.39 \pm 0.22$ |
| 8 | $0.78 \pm 0.11$ | $0.34 \pm 0.17$ |
| 7 | $0.80 \pm 0.10$ | $0.36 \pm 0.20$ |
| 6 | $0.78 \pm 0.11$ | $0.37 \pm 0.20$ |

We further compared the performance of SINCERITIES to three other network inference methods, namely TSNI [34], GENIE3 [36], and JUMP3 [35]. TSNI (Time Series Network Inference) is a GRN inference algorithm developed for time series gene expression data, relying on a linear ordinary differential equation model of the gene transcriptional process [34]. Meanwhile, GENIE3 (GEne Network Inference with Ensemble of trees) employs on a tree-based ensemble strategy using either random forest or extra-trees algorithms [36]. GENIE3 was among the top performers in DREAM 4 and DREAM 5 network inference challenges [44,45]. Recently, GENIE3 has also been applied to single cell data as a preliminary step to infer the skeleton of the GRN [31]. Lastly, JUMP3 uses a hybrid strategy combining non-parametric decision trees approach with dynamical ON/OFF modelling, to infer GRNs from time series expression data [35]. Since TSNI and JUMP3 require time series (longitudinal) data, we applied these methods to the (population) averages of the single cell gene expression data from each time point. Among the three previous methods, only TSNI generates GRN predictions with signed edges.

Figure 2 compares the AUROC and AUPR values of SINCERITIES and the three other methods mentioned above. The AUROC and AUPR values for TSNI and SINCERITIES were computed by respecting for the signs of the edges. However, for GENIE3 and JUMP3 predictions, the AUROC and AUPR values were based only on the existence of the regulatory edges (ignoring signs). The results showed that SINCERITIES significantly outperformed all of these methods (*p*-value < 0.05, paired t-tests) (see Additional file 2: Table S1).

**Figure 2.**
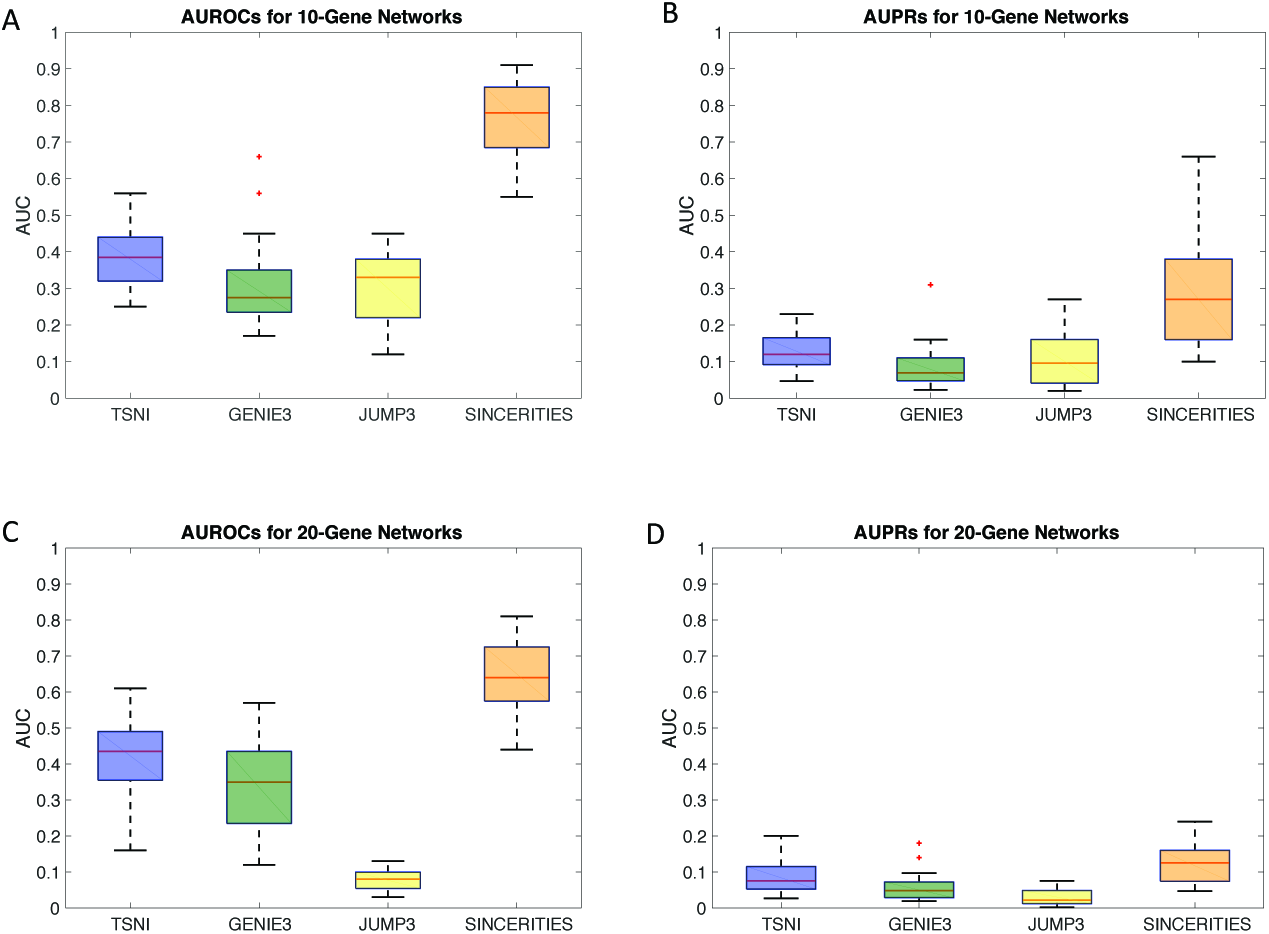
Performance comparison among TSNI, GENIE3, JUMP3, and SINCERITIES. (A) AUROC and (B) AUPR values for 10-gene gold standard GRNs. (C) AUROC and (D) AUPR values for 20-gene gold standard GRNs.

### Reconstructing GRN driving THP1 differentiation

In the following, we applied SINCERITIES to infer the GRN that drives the differentiation of monocytic THP-1 human myeloid leukemia cell differentiation into macrophages. The GRN of THP-1 differentiation has previously been constructed using deep sequencing (deepCAGE) and RNA interference (RNAi) experiments [46,47], providing the gold standard network for evaluating the performance of SINCERITIES and the three existing inference methods. The time-stamped cross-sectional single cell data came from qRT-PCR expression profiling of 45 transcription factors (TFs) in 960 THP-1 cells that were collected at 8 distinct time points (0, 1, 6, 12, 24, 48, 72, 96 hours) after stimulation by 12-myristate 13-acetate (PMA) [25].

We applied SINCERITIES as well as the three other methods to reconstruct the GRN of THP-1 differentiation using the single cell expression data above. The AUROC and AUPR values were evaluated using the anti-/pro-differentiation TF network found by RNAi knockdown experiments as the gold standard network [47]. We noted that only 20 TFs in the RNAi study overlapped with the set of genes in the single cell study [25]. Therefore, while the GRN inferences were done for 45 TFs, the calculation of AUROCs and AUPRs was based on the regulatory edges among the common set of 20 TFs. Again, for GENIE3 and JUMP3, the AUROC and AUPR values did not take into account the modes (signs) of the regulatory edges.

Table 3 shows the AUROCs and AUPRs for the four network inference strategies. For SINCERITIES, we reported the AUROC and AUPR values both with and without the mode of the gene regulations. The AUROC and AUPR values of SINCERITIES for the unsigned GRN prediction were similar to those using *in silico* data. As expected, the AUROC and AUPR values for signed GRN predictions were lower, but only slightly. TSNI, GENIE3, and JUMP3 performed worse than SINCERITIES, and did not give much better predictions than a random network (AUROC = 0.50).

**Table 3.** Performance comparison among TSNI, GENIE3, JUMP3 and SINCERITIES in inferring the GRN of THP-1 cell differentiation.

|  | **AUROC** | **AUPR** |
| --- | --- | --- |
| TSNI | 0.44 | 0.11 |
| GENIE3 | 0.46 | 0.23 |
| JUMP3 | 0.52 | 0.16 |
| SINCERITIES (without sign) | **0.70** | **0.33** |
| SINCERITIES (with sign) | **0.64** | **0.25** |

### Computational Runtime

To assess the computational complexity of our approach, we measured the runtimes of SINCERITIES for 10- and 20-gene *in silico* datasets, and compared these runtimes to those of the three other algorithms. Table 4 gives the average runtimes (in seconds) for TSNI, GENIE3, JUMP3 and SINCERITIES for the main *in silico* dataset and for the THP-1 differentiation data. Tree-based inference methods (GENIE3 and JUMP3) were significantly slower than SINCERITIES and TSNI. In particular, doubling the network size, the runtimes of GENIE3 and JUMP3 doubled and quadrupled, respectively. Meanwhile, the runtimes of SINCERITIES and TSNI finished almost instantaneously (<1 second) for these datasets, since these algorithms involved solving linear regressions. Finally, we noted that the regularized linear regressions in SINCERITIES (one for each gene) are independent of each other and are therefore amenable for parallel computation.

**Table 4.** Computational times comparison among TSNI, GENIE3, JUMP3 and SINCERITIES.

| **Average runtime (\*)** | **TSNI** | **GENIE3** | **JUMP3** | **SINCERITIES** |
| --- | --- | --- | --- | --- |
| 10-gene networks | 0.04 sec | 16 sec | 6 sec | 0.32 sec |
| 20-gene networks | 0.06 sec | 40 sec | 24 sec | 0.74 sec |
| THP-1 differentiation | 0.33 sec | 41 sec | 43 sec | 0.83 sec |

\* Computational times were measured on an 8-GB RAM, 1.6Ghz dual-core Intel core i5 computer.

### Comparison to other network inference methods for single cell data

The challenges of analyzing single cell transcriptional data have led to the creation of novel bioinfomatics algorithms, including algorithms for GRN inference using single cell transcriptional profiles [26–28,31]. A number of algorithms have been developed based on viewing the single cell gene expressions as binary state vectors, whose state transition trajectories are governed by a gene regulatory network with Boolean logic functions. Examples of such algorithms include SCNS [28], SingleCellNet [27] and BTR [29]. A general drawback of these algorithms is that the dimension of the state space of a Boolean network increases exponentially with respect to the number of genes (2^*m*^where *m* is the number of genes). Consequently, even for a moderately sized GRN (~50 genes), providing a reasonable coverage of the state space would require a tremendous number of single cell profiles. The extremely large state space will also make the inference problem computationally challenging.

Recently, Ocone et al. used a combination of a machine-learning algorithm GENIE3 and ODE modelling for GRN inference using single cell transcriptional data. Here, GENIE3 was first applied to produce a skeleton of the GRN. This skeleton was then refined by fitting an ODE model to pseudo-time trajectories of the gene expression, produced by applying Wanderlust algorithm [31] to single cell expression data in low-dimensional diffusion map projection [48]. However, there are several issues in using pseudo-time trajectories for GRN inference. First, one makes an implicit assumption that the trajectory reflects gene expression changes that are caused by the gene regulatory interactions associated with the physiological process of interest (e.g. cell differentiation). In our experience, the success of cell ordering in reproducing the gene expression trajectory depends sensitively on the cell sampling strategy, that is, being able to sample the right cells at the right time point or stages. For example, the application of Wanderlust to the *in silico* time-stamped single cell dataset from yeast led to cell ordering that was incongruent with the sampling times, especially for latter time points (Additional file 3: Fig. S1 and Fig. S2).

Meanwhile, Kouno et al. showed by using multiple dimension scaling that the THP-1 cell differentiation follows a rather irregular temporal dynamics in the low-dimensional (2D) projected space (see also Additional file 3: Fig. S3 for PCA, t-SNE and diffusion map analysis). As in the case of *in silico* dataset, Wanderlust ordering of THP-1 single cell expression data showed little correlation with the cell time-stamps (see Additional file 3: Fig. S4). SINCERITIES overcomes the issues mentioned above (large dataset requirement, high computational complexity, cell ordering) as the network inference involves numerically efficient regularized linear regression and directly use time-stamped cross-sectional data (without an intermediate step to construct state transition trajectory).

### Conclusion

Advances in single cell transcriptional profiling offer much promise in elucidating the functional role of cell-to-cell variability across different key physiological processes, such as stem cell differentiation. In particular, single cell expression data carry crucial information on the gene regulatory network that governs cellular heterogeneity and cell decision-making. However, inferring GRNs from single cell transcriptional profiles is complicated by the intrinsic stochasticity and bursty dynamics of the gene expression process and the loss of cell identity during high-throughput transcriptional profiling. In this work, we developed SINCERITIES for GRN inference using time-stamped cross-sectional single cell expression data, a common type of dataset generated by transcriptional profiling of single cells at multiple time points. SINCERITIES is based on the premise that changes in the gene expression distribution of a transcription factor in a given time window would cause a proportional change in the transcriptional expression distributions of the target genes in the next time window. The network inference involves numerically efficient ridge regression problem. In comparison to network inference algorithms for population average time series data (TSNI and JUMP3) and to a tree-based machine learning algorithm (GENIE3), SINCERITIES could provide significantly more accurate GRNs based on AUROCs and AUPRs.

## Methods

### Distribution Distance

In SINCERITIES, we used the Kolmogorov-Smirnov distance to quantify the distance between cumulative distribution functions of gene expressions from subsequent time points, according to

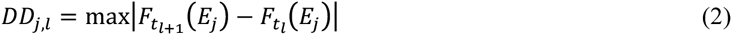

where *DD*_*j,l*_ denotes the distributional distance of gene *j* expression *E*_*j*_ between time points *t*_*l*_ and *t_l_+1* (l = 1, 2, …, *n—* 1) and *F*_*tl*_,(*E*_*j*_) denotes the cumulative distribution function of *E*_*j*_. We also evaluated two additional DD metrics, namely the Anderson-Darling (AD) statistics [49] and the Cramér-von Mises (CM) criterion [50] (see Additional file 3: Table S2). The performance of SINCERITIES did not depend sensitively on the DD metrics used. In order to accommodate non-uniformity in the sampling times, we normalized *DD*_*j,l*_ with respect to the time window size, as follows:

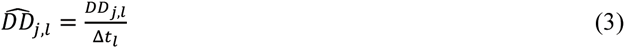

where 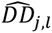 denotes the normalized distribution distance of gene *j* in the time window between *t*_*l*_ and *t*_*l+1*_ with △*t*_*l*_=*t*_*l+1*_—*t*_*l*_.

### Ridge Regression

As shown in Fig. 1C and in Eq. (1), for each gene *j*, we solved a linear regression problem of the form: **y** = **Xα**, where **y** denotes the *n*-2 vector of 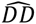distances of gene *j* corresponding to time windows Δ*t*_2_ to Δ*t*_*n*-1_,and **X** denotes the (*n*-2) x *m* matrix of 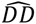 distances corresponding to time windows Δ*t*_1_ to Δ*t*_*n*-2_, for all genes. To obtain the solution vector **α**, we performed a ridge regression penalized least square optimization as follows:

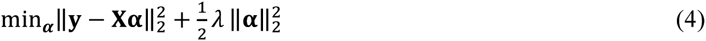

with the constraint that *α*_*i*_ ≥ 0. We used GLMNET algorithm (MATLAB) to generate the regularization path, i.e. the solution α as a function of different *λ* values [38].

The optimal weight factor *λ* is generally data dependent. Here, we performed a leave-one-out cross validation [51] to determine the optimal weight factor *λ*. In LOOCV, we allocated one row of **y** and **X** as the test dataset and the remaining as the training dataset. Then, we generated the regularization path for the training dataset using GLMNET, and computed the error of predicting the test dataset as a function of *λ*. We repeated this exercise for every permutation of test and training dataset assignment, and selected the optimal λ that minimized the average prediction error. Finally, we ran GLMNET on the full dataset and took the solution *α*_*_ that corresponded to the optimal *λ* value above. In addition to ridge regression, we also tested SINCERITIES with two other penalty functions: the ‘Least Absolute Shrinkage and Selection Operator’ (Lasso) L1-norm penalty [52], which was adopted in our previous algorithm SNIFS (Sparse Network Inference For Single cell data) [53], and the elastic-net penalty [54]. These alternative penalty functions however led to much poorer GRN predictions than ridge regression (for further details, see additional file 3: Table S3).

### Partial Correlation Analysis

In order to determine the mode (sign) of gene regulatory relationships, we performed the Spearman rank partial correlation analysis. More specifically, for each time point *k* (*k* = 1, 2, …, *n*), we calculated the Spearman rank partial correlation coefficient of gene expressions from every pair of genes while controlling for the other genes. The sign of the regulatory edge pointing from gene *i* to gene *j* was set equal to the sign of the partial correlation coefficient for the combined expression data over all time points. Note that by using correlation, the sign of the edge pointing from gene *i* to gene *j* is equal to the sign of the edge pointing from gene *j* to gene *i*.

### In silico data generation

We used GeneNetWeaver (GNW) to randomly generate 10-gene and 20-gene random subnetworks of *Escherichia coli* and *Saccharomyces cerevisiae* (yeast) GRNs. After removing self-regulations, we simulated *in silico* single cell expression data using the following stochastic differential equation (SDE) model of the mRNA [55]:

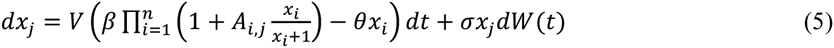

where *x*_*j*_ describes the mRNA level of gene *j*, *A*_*i,j*_ denotes the regulation of the expression of gene *j* by gene *i*, *β* denotes the basal transcriptional rate, *θ* denotes the mRNA degradation rate constant, and *σ* and *V* are scaling parameters. The term *dW(t)* denotes the random Wiener process, simulating the intrinsic stochastic dynamics of gene expression [56]. We set *A*_*ij*_ to 1 for gene activation, to −1 for gene repression, and to 0 otherwise. For the main dataset in the case study, we set the parameters to the following: *V*=30, *β*=1, θ=0.2, and *σ*=0.1.

We simulated the SDE model above using the Euler-Maruyama method [57] with an initial condition *x*_*j*_(0) set to 0 for every gene, until the gene expression reached steady state (*t* = 3 arbitrary time unit). For each GRN structure, we generated 100 stochastic trajectories for each time point (a total of 8×100 =800 independent trajectories for 8 time points), representing 100 single cells. The simulations above mimicked the scenario where single cells are lysed for gene expression profiling [28,58]. To test the robustness of SINCERITIES with respect to the intrinsic noise in gene expression and to the number of sampling time points, we further generated two supplementary datasets from the 20 10-gene *E.coli* and yeast gold standard GRNs: by varying *σ* parameter between 0.1 and 0.4 with a step of 0.1 (see Table 2A) and by selecting the first *n* time points from the following set *t* = 0.51, 0.60, 0.74, 1.2, 1.3, 1.5, 1.8, 2.2, 2.6, and 3 where *n* is between 6 and 10. The time points had been selected to exclude the time period when the mRNA level rose quickly from the initial concentration. This initial increase was a consequence of starting the simulations from *x*_*j*_ (0) = 0, and did not necessarily reflect the gene regulatory actions.

## Software availability

The MATLAB implementation of SINCERITIES is freely available from the following website: http://www.cabsel.ethz.ch/tools/sincerities.html.

## Availability of data and materials

The single cell THP-1 transcriptional profiles are available from the original publications (see supplementary material in [25]). The *in silico* single cell data are available from SINCERITIES website: http://www.cabsel.ethz.ch/tools/sincerities.html">.

## Ethics approval

Ethics approval is not applicable for this study.

## Additional file

Additional file 1: The gold standard GRN of *E.coli* and yeast used for the generation of single cell *in silico* gene expression data.

Additional file 2: Table S1. Performance evaluation for TSNI, GENIE3, JUMP3, and SINCERITIES with Ridge regression and KS distance.

Additional file 3: Supplementary material.

## Acknowledgements

We would like to thank Dr. Rajanikanth Vadigepali (Thomas Jefferson University) for fruitful discussions, and Ms. Heeju Noh for assistance in MATLAB implementation.

## References

1. Pieprzyk M, High H. Fluidigm Dynamic Arrays provide a platform for single-cell gene expression analysis. Nat. Methods. 2009;6.

2. Gupta PB, Fillmore CM, Jiang G, Shapira SD, Tao K, Kuperwasser C, et al. Stochastic state transitions give rise to phenotypic equilibrium in populations of cancer cells. Cell. 2011;146:633–44.

3. Shalek AK, Satija R, Shuga J, Trombetta JJ, Gennert D, Lu D, et al. Single-cell RNA-seq reveals dynamic paracrine control of cellular variation. Nature. 2014;510.

4. Pollen AA, Nowakowski TJ, Shuga J, Wang X, Leyrat AA, Lui JH, et al. Low-coverage single-cell mRNA sequencing reveals cellular heterogeneity and activated signaling pathways in developing cerebral cortex. Nat. Biotechnol. 2014;32:1053–8.

5. Kumar RM, Cahan P, Shalek AK, Satija R, DaleyKeyser AJ, Li H, et al. Deconstructing transcriptional heterogeneity in pluripotent stem cells. Nature. 2014;516:56–61.

6. Buettner F, Natarajan KN, Casale FP, Proserpio V, Scialdone A, Theis FJ, et al. Computational analysis of cell-to-cell heterogeneity in single-cell RNA-sequencing data reveals hidden subpopulations of cells. Nat. Biotechnol. 2015;33:155–60.

7. Chang HH, Hemberg M, Barahona M, Ingber DE, Huang S. Transcriptome-wide noise controls lineage choice in mammalian progenitor cells. Nature. 2008;453:544–7.

8. Fang M, Xie H, Dougan SK, Ploegh H, van Oudenaarden A. Stochastic cytokine expression induces mixed T helper cell States. PLoS Biol. 2013;11.

9. Lee J, Lee J, Farquhar KS, Yun J, Frankenberger CA, Bevilacqua E, et al. Network of mutually repressive metastasis regulators can promote cell heterogeneity and metastatic transitions. Proc. Natl. Acad. Sci. U. S. A. 2014;111:364–73.

10. Kim K-T, Lee HW, Lee H-O, Kim SC, Seo YJ, Chung W, et al. Single-cell mRNA sequencing identifies subclonal heterogeneity in anti-cancer drug responses of lung adenocarcinoma cells. Genome Biol. 2015;16:127.

11. Richard A, Boullu L, Herbach U, Bonnafoux A, Morin V, Vallin E, et al. A surge in cell-to-cell molecular variability precedes the commitment in a differentiation process. PLoS Biol. In press.

12. Simpson EH. The Interpretation of Interaction in Contingency Tables. J. R. Stat. Soc. Ser. B. 1951;13:238–41.

13. Liu S, Trapnell C. Single-cell transcriptome sequencing: recent advances and remaining challenges. F1000Research. 2016;5.

14. Stegle O, Teichmann SA, Marioni JC. Computational and analytical challenges in single-cell transcriptomics. Nat. Rev. Genet. 2015;16:133–45.

15. Bacher R, Kendziorski C, Auer P, Doerge R, Robles J, Qureshi S, et al. Design and computational analysis of single-cell RNA-sequencing experiments. Genome Biol. 2016;17:63.

16. Amir ED, Davis KL, Tadmor MD, Simonds EF, Levine JH, Bendall SC, et al. viSNE enables visualization of high dimensional single-cell data and reveals phenotypic heterogeneity of leukemia. Nat. Biotechnol. 2013;31:545–52.

17. Buettner F, Moignard V, Göttgens B, Theis FJ. Probabilistic PCA of censored data: accounting for uncertainties in the visualization of high-throughput single-cell qPCR data. Bioinformatics. 2014;30:1867–75.

18. Haghverdi L, Buettner F, Theis FJ. Diffusion maps for high-dimensional single-cell analysis of differentiation data. Bioinformatics. 2015;2989–98.

19. Xu C, Su Z. Identification of cell types from single-cell transcriptomes using a novel clustering method. Bioinformatics. 2015;31:1974–80.

20. Pierson E, Yau C, Shapiro E, Biezuner T, Linnarsson S, Blainey P, et al. ZIFA: Dimensionality reduction for zero-inflated single-cell gene expression analysis. Genome Biol. 2015;16:241.

21. Trapnell C, Cacchiarelli D, Grimsby J, Pokharel P, Li S, Morse M, et al. The dynamics and regulators of cell fate decisions are revealed by pseudotemporal ordering of single cells. Nat. Biotechnol. 2014;32:381–6.

22. Bendall SC, Davis KL, Amir E-AD, Tadmor MD, Simonds EF, Chen TJ, et al. Single-cell trajectory detection uncovers progression and regulatory coordination in human B cell development. Cell. 2014;157:714–25.

23. Marco E, Karp RL, Guo G, Robson P, Hart AH, Trippa L, et al. Bifurcation analysis of single-cell gene expression data reveals epigenetic landscape. Proc. Natl. Acad. Sci. U. S. A. 2014;111:5643–50.

24. Ji Z, Ji H. TSCAN: Pseudo-time reconstruction and evaluation in single-cell RNA-seq analysis. Nucleic Acids Res. 2016;117.

25. Kouno T, de Hoon M, Mar JC, Tomaru Y, Kawano M, Carninci P, et al. Temporal dynamics and transcriptional control using single-cell gene expression analysis. Genome Biol. 2013;14:R118.

26. Pina C, Teles J, Fugazza C, May G, Wang D, Guo Y, et al. Single-Cell Network Analysis Identifies DDIT3 as a Nodal Lineage Regulator in Hematopoiesis. Cell Rep. 2015;11:1503–10.

27. Chen H, Guo J, Mishra SK, Robson P, Niranjan M, Zheng J. Single-cell transcriptional analysis to uncover regulatory circuits driving cell fate decisions in early mouse development. Bioinformatics. 2014;31:1060–6.

28. Moignard V, Woodhouse S, Haghverdi L, Lilly AJ, Tanaka Y, Wilkinson AC, et al. Decoding the regulatory network of early blood development from single-cell gene expression measurements. Nat. Biotechnol. 2015;33:269–76.

29. Lim CY, Wang H, Woodhouse S, Piterman N, Wernisch L, Fisher J, et al. BTR: training asynchronous Boolean models using single-cell expression data. BMC Bioinformatics. 2016;17:355.

30. Teles J, Pina C, Edén P, Ohlsson M, Enver T, Peterson C. Transcriptional regulation of lineage commitment–a stochastic model of cell fate decisions. PLoS Comput. Biol. 2013;9:e1003197.

31. Ocone a., Haghverdi L, Mueller NS, Theis FJ. Reconstructing gene regulatory dynamics from high-dimensional single-cell snapshot data. Bioinformatics. 2015;31:i89–96.

32. Bar-Joseph Z, Gitter A, Simon I. Studying and modelling dynamic biological processes using time-series gene expression data. Nat. Rev. Genet. 2012;13:552–64.

33. Chu L-F, Leng N, Zhang J, Hou Z, Mamott D, Vereide DT, et al. Single-cell RNA-seq reveals novel regulators of human embryonic stem cell differentiation to definitive endoderm. Genome Biol. 2016;17:173.

34. Bansal M, Gatta G Della, Di Bernardo D. Inference of gene regulatory networks and compound mode of action from time course gene expression profiles. Bioinformatics. 2006;22:815–22.

35. Huynh-Thu VA, Sanguinetti G. Combining tree-based and dynamical systems for the inference of gene regulatory networks. Bioinformatics. 2015;31:1614–22.

36. Huynh-Thu VA, Irrthum A, Wehenkel L, Geurts P. Inferring regulatory networks from expression data using tree-based methods. PLoS One. 2010;5:e12776.

37. Massey FJ. The Kolmogorov-Smirnov Test for Goodness of Fit. J. Am. Stat. Assoc. 1951;46:68–78.

38. Friedman J, Hastie T, Tibshirani R. Regularization Paths for Generalized Linear Models via Coordinate Descent. J. Stat. Softw. 2010;33:1–22.

39. Huang W, Cao X, Biase FH, Yu P, Zhong S. Time-variant clustering model for understanding cell fate decisions. Proc. Natl. Acad. Sci. U. S. A. 2014;111:E4797–806.

40. Szederkényi G, Banga JR, Alonso AA. Inference of complex biological networks: distinguishability issues and optimization-based solutions. BMC Syst. Biol. 2011;5:177.

41. Ud-Dean SMM, Gunawan R. Ensemble inference and inferability of gene regulatory networks. PLoS One. 2014;9:e103812.

42. Schaffter T, Marbach D, Floreano D. GeneNetWeaver: in silico benchmark generation and performance profiling of network inference methods. Bioinformatics. 2011;27:2263–70.

43. Marbach D, Prill RJ, Schaffter T, Mattiussi C, Floreano D, Stolovitzky G. Revealing strengths and weaknesses of methods for gene network inference. Proc. Natl. Acad. Sci. U. S. A. 2010;107:6286–91.

44. Marbach D, Schaffter T, Mattiussi C, Floreano D. Generating realistic in silico gene networks for performance assessment of reverse engineering methods. J. Comput. Biol. 2009;16:229–39.

45. Marbach D, Costello JC, Küffner R, Vega NM, Prill RJ, Camacho DM, et al. Wisdom of crowds for robust gene network inference. Nat. Methods. 2012;9:796–804.

46. Vitezic M, Lassmann T, Forrest ARR, Suzuki M, Tomaru Y, Kawai J, et al. Building promoter aware transcriptional regulatory networks using siRNA perturbation and deepCAGE. Nucleic Acids Res. 2010;38:8141–8.

47. Tomaru Y, Simon C, Forrest AR, Miura H, Kubosaki A, Hayashizaki Y, et al. Regulatory interdependence of myeloid transcription factors revealed by Matrix RNAi analysis. Genome Biol. 2009;10:R121.

48. Coifman RR, Lafon S. Diffusion maps. Appl. Comput. Harmon. Anal. 2006;21:5–30.

49. Anderson TW, Darling DA. Asymptotic Theory of Certain “Goodness of Fit” Criteria Based on Stochastic Processes. Ann. Math. Stat. Institute of Mathematical Statistics; 1952;23:193–212.

50. Anderson TW. On the Distribution of the Two-Sample Cramer-von Mises Criterion. Ann. Math. Stat. 1962;33:1148–59.

51. Kohavi R. A study of cross-validation and bootstrap for accuracy estimation and model selection. 1995;1137–43.

52. Tibshirani R. Regression shrinkage and selection via the lasso. J. R. Stat. Soc. Ser. B. 1996. p. 267–88.

53. Papili Gao N, Ud-dean SMM, Gunawan R. Gene Regulatory Network Inference Using Time-Stamped Cross-Sectional Single Cell Expression Data. Proc. Found. Syst. Biol. Eng. 2016. In press.

54. Zou H, Hastie T. Regularization and variable selection via the elastic net. J. R. Stat. Soc. Ser. B. 2005;67:301–20.

55. Pinna A, Soranzo N, de la Fuente A. From knockouts to networks: establishing direct cause-effect relationships through graph analysis. PLoS One. 2010;5:e12912.

56. Wilkinson DJ. Stochastic modelling for quantitative description of heterogeneous biological systems. Nat. Rev. Genet. Nature Publishing Group; 2009;10:122–33.

57. Higham. DJ. An Algorithmic Introduction to Numerical Simulation of Stochastic Differential Equations. SIAM Rev. Society for Industrial and Applied Mathematics; 2001;43:525–46.

58. Gao W, Zhang W, Meldrum DR. RT-qPCR based quantitative analysis of gene expression in single bacterial cells. J. Microbiol. Methods. 2011;85:221–7.

